# Quantification of RNA Polymerase I transcriptional attenuation at the active VSG expression site in *Trypanosoma brucei*

**DOI:** 10.1101/2021.06.21.449234

**Authors:** Nadine Weisert, Klara Thein, Helena Reis, Christian J. Janzen

**Author notes:** To whom correspondence should be addressed. Christian J. Janzen.

## Abstract

The cell surface of the extracellular pathogen *Trypanosoma brucei* consists of a dense coat of variant surface glycoprotein (VSG), which enables the parasite to evade the immune system of the vertebrate host. Only one VSG gene from a large repertoire is expressed from a so-called bloodstream form expression site (BES) at a given timepoint. There are several BES in every parasite but only one is transcriptionally active. Other BES are silenced by transcriptional attenuation. Periodic activation of a previously-silenced BES results in differential VSG transcription and escape from the immune response. A process called antigenic variation. In contrast to gene transcription in other eukaryotes, the BES is transcribed by RNA polymerase I (Pol I). It was proposed that this highly-processive polymerase is needed to provide a sufficiently high transcription rate at the VSG gene. Surprisingly, we discovered a position-dependent Pol I activity and attenuation of transcriptional elongation also at the active BES. Transcription rates at the VSG gene appear to be comparable to Pol II-mediated transcription of house-keeping genes. Although these findings are in contradiction to the long-standing concept of continuously high transcription rates at the active BES in *Trypanosoma brucei*, they are complementary to recent groundbreaking findings about transcriptional regulation of VSG genes.

## INTRODUCTION

The unicellular parasite *Trypanosoma brucei*, which belongs to the protist class of Kinetoplastea, is the causative agent of the African trypanosomiasis [1]. *T. brucei* alternates between two main life cycle stages, the bloodstream form (BSF) in the mammalian host and the procyclic form (PCF) in the tsetse fly vector [2]. In the mammalian host, the parasite evades the immune system by alternately expressing a different copy of variant surface glycoprotein (VSG) genes [3, 4]. This antigenic variation enables the pathogen to survive in the extracellular environment and causes chronic infections [5]. The transcription of *VSG* genes takes place from one of several so-called bloodstream form expression sites (ES), which are polycistronic transcription units located adjacent to telomeres. BES are composed of a promoter, a variable number of expression site-associated genes (ESAGs), 70 bp tandem repeats and the VSG gene, located proximal to the telomeric repeat region [6]. The entire BES can span up to 15 kilobase pairs. Only one BES is fully active at any given timepoint and therefore only one VSG variant is expressed, while all other BES are silenced by transcriptional attenuation. Changing of the VSG coat is mediated either by *in situ* or recombinational switching, using the large repertoire of over 2000 VSG genes, pseudogenes and gene fragments [7, 8].

An unusual feature of the BES is that transcription is mediated by RNA Polymerase I (Pol I), which typically only transcribes ribosomal RNA genes in other eukaryotes [9]. In trypanosomes, Pol I transcribes not only VSG genes and rRNA genes but is also responsible for transcription of procyclin mRNA, which encodes for the surface protein in insect stage trypanosomes [10]. This is possible due to another trypanosome-specific mechanism, trans-splicing. Usually, mRNA processing - such as 5’-capping - is mediated by interaction of the c-terminal subunit of RNA polymerase II (Pol II) complex with capping enzymes. In trypanosomes, capping and transcription are not coupled. A so-called splice leader sequence is added post-transcriptionally, which enables Pol I to produce functional mRNAs [11, 12]. *T. brucei* has a high demand for VSG mRNA in the BSF in order to establish the densely-packed surface coat. VSG protein represents approximately 10% of total protein mass in BSF [13]. Due to the fact that expression takes place from a single gene locus, Pol I transcription of VSG is proposed to be substantially higher than Pol II transcription of housekeeping genes like tubulin [14, 15].

Different processes, such as regulation of promoter activity, changes of chromatin structure and effects of telomere-associated protein complexes influence VSG expression. For example, it has been proposed that post-translational modifications regulate BES transcription on an epigenetic level by altering chromatin organization and therefore DNA accessibility [16]. The active BES has an open, nucleosome-depleted chromatin structure while silent BES are nucleosome-associated [17]. The chromatin remodeling complex ISWI, the histone methyltransferase DOT1B, histones H1 and H3 and histone chaperons such as chromatin assembly factor-1 (CAF-1b), FACT and anti-silencing factor 1A (ASF1A) are all involved in the maintenance of monoallelic expression of the active BES [18-22]. For example, depletion of the linker histone H1 results in 6-fold higher transcriptional rate at promoter proximal regions in silent BES [20, 23]. The Class I transcription factor A (CITFA) is important for transcription initiation and the trypanosome DNA binding protein 1 (TDP1) is involved in nucleosome depletion at the active BES [24, 25]. Furthermore, the active BES is associated with an extranuclear expression site body (ESB), containing Pol I foci [26]. Recently, several proteins such as VSG exclusion 1 and 2 and CAF-1 have been identified to be associated with the ESB and are important to establish and control monoallelic exclusion [27, 28]. Several reports describe the involvement of telomere-binding or -associating proteins in VSG regulation. For example, TTAGGG repeat factor (TRF) and TRF-interacting factor 2 (TIF2) suppress VSG switching, repressor/activator protein 1 (RAP1) is important to maintain monoallelic expression and telomere-associated protein 1 (TelAP1) regulates developmental transcriptional silencing [29-32]. During the developmental transition to the PCF in the tsetse fly vector, all VSG transcription is silenced and the transcription level of all BES is reduced to 2% [25]. The ESB dissolves while a migration of the active BES to the nuclear periphery and chromatin reorganization takes place [33]. Using luciferase reporter cassettes integrated at different positions in an BES, Navarro *et al*. proposed that developmental silencing is accompanied by chromatin condensation which leads to reduced accessibility of the entire BES [34]. Silencing of a BES in BSF appears to be mediated by a different mechanism. Most likely, Pol I is installed and active at the promotor of all BES [35]. Promoter proximal transcription results in 20 % of ESAG mRNA originating from silent BES. Transcriptional processivity, however, is attenuated in silent BES and the Pol I complex does not reach the VSG gene at the end of the BES. It is assumed that in an active BES, the RNA elongation/processing machinery allows high transcriptional activity without attenuation throughout the entire BES.

In previous work however, we observed that luciferase reporter cassettes inserted immediately downstream of the promoter or proximal to the telomere resulted in substantially different luciferase activity readings in an active BES [36]. To further analyze these observations, we integrated a Renilla luciferase reporter either near the promoter region or proximal to the telomere. In both cell lines, luciferase mRNA processing was mediated by identical untranslated regions for better resemblance and comparability of luciferases activity. Unexpectedly, we also observed a partial attenuation of transcriptional elongation at the active BES, comparable with observations thus far only shown for silent BES. This stands in contradiction to previous assumptions regarding the transcriptional activity of the active BES.

## MATERIALS AND METHODS

### Trypanosome cell line and cultivation

Cell lines used in this study were based on monomorphic *Trypanosoma brucei* single marker (SM) bloodstream form (Lister strain 427 antigenic type MITat 1.2, clone 221a, modified with pHD328 and pLew114hyg5’) [37] or the 2T1 bloodstream form (Lister strain 427 antigenic type MITat 1.2, clone 221a, modified with pHD1313 and ph3Ep) [38]. Cells were cultured in HMI-9 medium at 37°C and 5% CO_2_ containing 10% heat-inactivated fetal calf serum [39]. Stable transfection and drug selection were carried out as described [40]. Cell population density was measured using Beckman Coulter^™^ Z2 particle count and size analyzer.

### Plasmid construction and transgenic cell lines

To generate luciferase reporter cell lines, the SM or 2T1 cell line was transfected with either linearized plasmids or with PCR-amplified fragments. The concentration of antibiotics was 5-10 times higher than normally recommended to confirm integration in the active BES [19]. For integration of luciferase reporter constructs in the active BES downstream of the promoter, a region between the BES transcription start site and ESAG7 was chosen. For integration of constructs near the telomere, the telomere-proximal Ψ221 pseudogene region was used as the integration site. Exact sequences can be provided upon request.

#### SR-I

SM cells were transfected with BstApI-linearized pCJ25A-Rluc. The plasmid was generated using pCJ25A by replacing the Firefly luciferase gene with the Renilla luciferase gene from pFG14 [19, 41]. pCJ25A-Rluc consists of a homologous region of Ψ221 for telomere-proximal integration. The Renilla luciferase is flanked by tubulin untranslated regions (UTRs), the blasticidin S deaminase gene (BSD) is flanked by aldolase UTRs.

#### SR-II

SM cells were transfected with linearized pFG14n (digested with KpnI/SacI). pFG14n consist of a homologous region for integration downstream of the BES promoter and a Renilla luciferase flanked by tubulin UTRs. The hygromycin phosphotransferase gene is flanked by actin UTRs.

#### DR-I: Fluc (pro) Rluc (tel)

SM cells were transfected with linearized pPur-Luc (digested with KpnI/NotI) [31] and pCJ25A-Rluc. The pPur-Luc plasmid consists of a homologous region for integration downstream of the BES promoter and a Firefly luciferase flanked by a 5’GPEET UTR and a 3’aldolase UTR. The puromycin N-acetyl-transferase gene (PUR) is flanked by aldolase UTRs.

#### DR-II: Rluc (pro) Fluc (tel)

SM cells were transfected with BstApI-linearized pFG14n and pCJ25A. pCJ25A consists of a homologous region for telomere-proximal integration in Ψ221 and a Firefly luciferase flanked by a 5’actin and a 3’aldolase UTR.

#### P10 DR-II

This cell line was provided by L. Figueiredo (University of Lisbon) [19]. PUR was integrated downstream of BES promoter in active BES1 (pLF12), the aminoglycoside phosphotransferase resistance gene (NEO) was integrated downstream of BES promoter in silent BES17 (pFL13). Luciferase integration occurred by transfection with linearized pCJ25A and pFG14n, replacing pLF12. The P10 DR-II switcher cell line was obtained by selection with G418 (active BES17).

##### Pol II-Rluc

2T1 cells were transfected with a PCR-amplified fragment derived from pFG14-NEO. pFG14 contains a Renilla luciferase gene flanked by tubulin UTRs followed by an aminoglycoside phosphotransferase resistance gene (NEO) flanked by actin UTRs. The primers for amplification contained 80bp of homologous regions of the alpha tubulin locus.

##### Pol II-Fluc

SM cells were transfected with linearized pCW37 (digested with NotI/XhoI) [42]. pCW37 consists of a homologous region for integration into a locus within a PTU on chromosome 1, a firefly luciferase flanked by a 5’GPEET UTR and a 3’aldolase UTR. The hygromycin phosphotransferase gene is flanked by actin UTRs.

### Differentiation of BSF into PCF

For differentiation of monomorphic BSF, cells were grown to a cell density of 1.5 ⨯ 10^6^ cells/ml. A total of 3 ⨯ 10^7^ cells were harvested by centrifugation (1500 x g, 10 min, RT) and the cell pellet was resuspended in 5 ml DTM medium containing 6 mM cis-aconitate [43]. Cells were then cultured at 27°C and 5% CO_2_. SDM-79 medium containing 10% heat-inactivated fetal calf serum was used for further dilution [44].

### Dual-luciferase assay

Renilla luciferase (Rluc) and Firefly luciferase (Fluc) activity was measured using the dual-luciferase reporter assay kit (Promega). Measurements were performed according to the manufacturer’s instructions, as described previously [32]. Briefly, 1 × 10^6^ cells were harvested by centrifugation, washed with ice-cold phosphate-buffered saline (PBS) before further centrifugation (1500 x g, 10 min, 4°C) and then resuspended in 100 µl 1 x Passive Lysis Buffer (PLB, Promega). 45 µl of the luciferase assay substrate dissolved in Luciferase Assay Buffer II was plated on a 96-well plate and 10 µl of the cell lysate was added immediately before measurement of Fluc activity using a Tecan reader. 45µl of Stop & Glo Reagent was added to quench the Fluc luminescence before the Rluc activity was measured directly afterwards.

### Western blot analysis

Whole cell lysates from 1 ⨯ 10^6^ cells were separated on 10% SDS-PAGE gels and transferred onto PVDF membranes. Blocking of the membrane was done in 5% milk powder, dissolved in PBS. The membrane was incubated with antibodies in PBS/0.1 % Tween 20 solution for 1h at RT. After each antibody incubation three washes with PBS/0.2 % Tween 20 solution followed. The CRD-depleted anti-VSG-2 (concentration 1:10,000) as well as anti-VSG-13 (concentration 1:100,000) rabbit antibodies were provided by L. Figueiredo (University of Lisbon) [19], the monoclonal mouse anti-PFR1,2 antibody L13D6 (concentration 1:500) was a gift from K. Gull (University of Oxford)[45]. Primary antibodies were detected using IRdye 680- and 800- coupled antibodies.

## RESULTS

### Monitoring of locus-specific differences in transcriptional activity during differentiation

Our previous work suggested that the kinetics of transcriptional silencing of reporter genes in the BES might be different depending on the location within the BES [32]. However, due to the experimental design of these studies, the signals from the reporter genes used were not directly comparable. We therefore decided to improve the reporter cell lines to evaluate transcriptional regulation of an active BES at different loci during *in situ* switching and differentiation in a quantitative manner. To monitor the transcriptional silencing process of the active BES during developmental differentiation, we generated two novel luciferase-based reporter cell lines. We integrated a Renilla luciferase (Rluc) reporter gene at one of two different loci in the expression site 1 (ES1). One reporter was located at a telomere proximal region, integrated in the VSG pseudogene Ψ221 (single reporter cell line I, SR-I). The other reporter was integrated into a region between the promoter and ESAG7, creating the SR-II cell line (**Error! Reference source not found**.A). Since the structure of the reporter cassettes are identical, we assumed that luciferase activity reflects RNA polymerase activity at the individual locus [46]. Integration into the active BES was ensured by using up to 10-fold higher antibiotic concentrations compared to standard selection for integration into chromosome internal loci [19]. Maintenance of VSG-2 expression was confirmed by Western blot analysis (Supplementary Fig S1). The Rluc is flanked by tubulin UTR’s, facilitating mRNA processing without influencing mRNA stability during differentiation [47]. After integration of the same luciferase reporter genes with identical UTRs, we could directly compare the expression intensity. After induction of differentiation, luciferase activity was measured at defined timepoints for four days (**Error! Reference source not found**.B). For compensation of background signals, the SM cell line without reporters was included and relative light units were deducted from the reporter activity. The luciferase reporter signal at the telomeric region increased 1.3-fold immediately after induction of differentiation, but already displayed approximately half of the activity 48 h post-induction. These results confirmed and quantified the observations made previously [32] and indicated faster BES silencing kinetics at the telomeric region compared to promoter proximal sites. Surprisingly, the absolute values of the Rluc signals revealed that the promoter luciferase activity was more than 70-fold higher at 0h compared to the luciferase activity at the telomeric region (**Error! Reference source not found**.C). This difference increased to more than 200-fold 48h post induction of differentiation (Supplementary Fig S2). These substantial differences were unexpected and indicated the active BES is also affected by transcriptional attenuation.

### Transcriptional activity at the active BES points to partial attenuation of transcriptional elongation

To confirm those results and to exclude a luciferase-dependent effect, we generated two dual reporter cell lines. In the first cell line we integrated a Firefly luciferase (Fluc) reporter gene at the promoter region and Rluc at the telomere region (DR-I) while the second reporter cell line was constructed vice versa (DR-II) as described in Reis et al (**Error! Reference source not found**.A) [32]. The luciferase activities of these cell lines could now be compared directly (**Error! Reference source not found**.B). Here, a 75-fold higher activity of the Rluc reporter and a 16-fold higher activity of the Fluc reporter was observed at the promoter integration sites compared to the telomere proximal locus. DR cell lines were also subjected to developmental differentiation and luciferase activity was measured at defined time points for 8 days, showing similar results as the SR (**Error! Reference source not found**.C vs. **Error! Reference source not found**.B). To confirm that all cell lines were still expressing VSG-2 and did not switch, whole cell lysates were analyzed by Western blot (Supplementary Fig S1). To confirm that integration of the reporter constructs did not impair transcriptional regulation or switching capacity of the BES, DR-II constructs were integrated into the active BES (BES1) of the P10 cell line [19]. With this cell line, it is possible to select for *in situ* switching to the previous silent BES17 using an aminoglycoside phosphotransferase resistance gene, which is integrated downstream of the BES17 promoter (**Error! Reference source not found**.A). Measurements of luciferase expression in active and silent BES could now be compared to verify accurate silencing after an *in situ* switch (**Error! Reference source not found**.B). Luciferase reporters in P10 DR-II in the active BES1 showed similar activity compared to DR-II. After switching, more than 100-fold reduction of the Rluc signal from the promoter region could be observed whereas the Fluc signal dropped to background signal values suggesting that transcriptional regulation during an *in situ* switch is comparable to wild-type parasites. To confirm a successful switching event, whole cell lysates were analyzed by Western blot techniques using VSG-2- and VSG-13-specific antibodies (Supplementary Fig S3). SM (expressing VSG-2) and N50 (expressing VSG-13) whole cell lysates served as controls. The switcher cell line was also switched back to the formerly active BES (BES1) to exclude recombination events in the corresponding BESs of the system. Whole cell lysates of this cell line are included in Western blot analysis shown in Fig S3. In summary, these data confirmed functional integration of the luciferase reporters in the active BES of the reporter cell line and suggests a substantial transcriptional attenuation of active BES in trypanosomes.

### The Pol I transcriptional activity at the telomere proximal region of the active BES is comparable to Pol II activity

Pol I-mediated transcription initiation is ten-fold higher compared to Pol II-driven transcription initiation of a PTU [14, 15]. However, to our knowledge, activity of luciferase reporter genes integrated in BES adjacent to telomeres have never be quantitatively compared with reporters integrated into PTUs. To address this question, two reporter cell lines were constructed. One contained the Fluc gene flanked by a 5’GPEET UTR and a 3’aldolase UTR integrated into a PTU (Pol II-Fluc) as described previously [42]. The other cell line contained a Rluc gene flanked by tubulin UTRs integrated within an array of tubulin genes (Pol II-Rluc) (**Error! Reference source not found**.A). The Fluc and Rluc signal intensities were compared with those of the DR-II and DR-I signals (Fig 2A), respectively. As describe already, we could detect a substantial difference between BES promoter-proximal reporter activity and Fluc and Rluc signal intensities transcribed from PTUs (Fig 4B). However, there was no substantial difference between PTU-derived reporter activity and signals from telomeric luciferase genes, indicating that the transcriptional activity of Pol II in PTUs is similar to that of Pol I at a telomere proximal region of the active BES.

**Fig 1:**
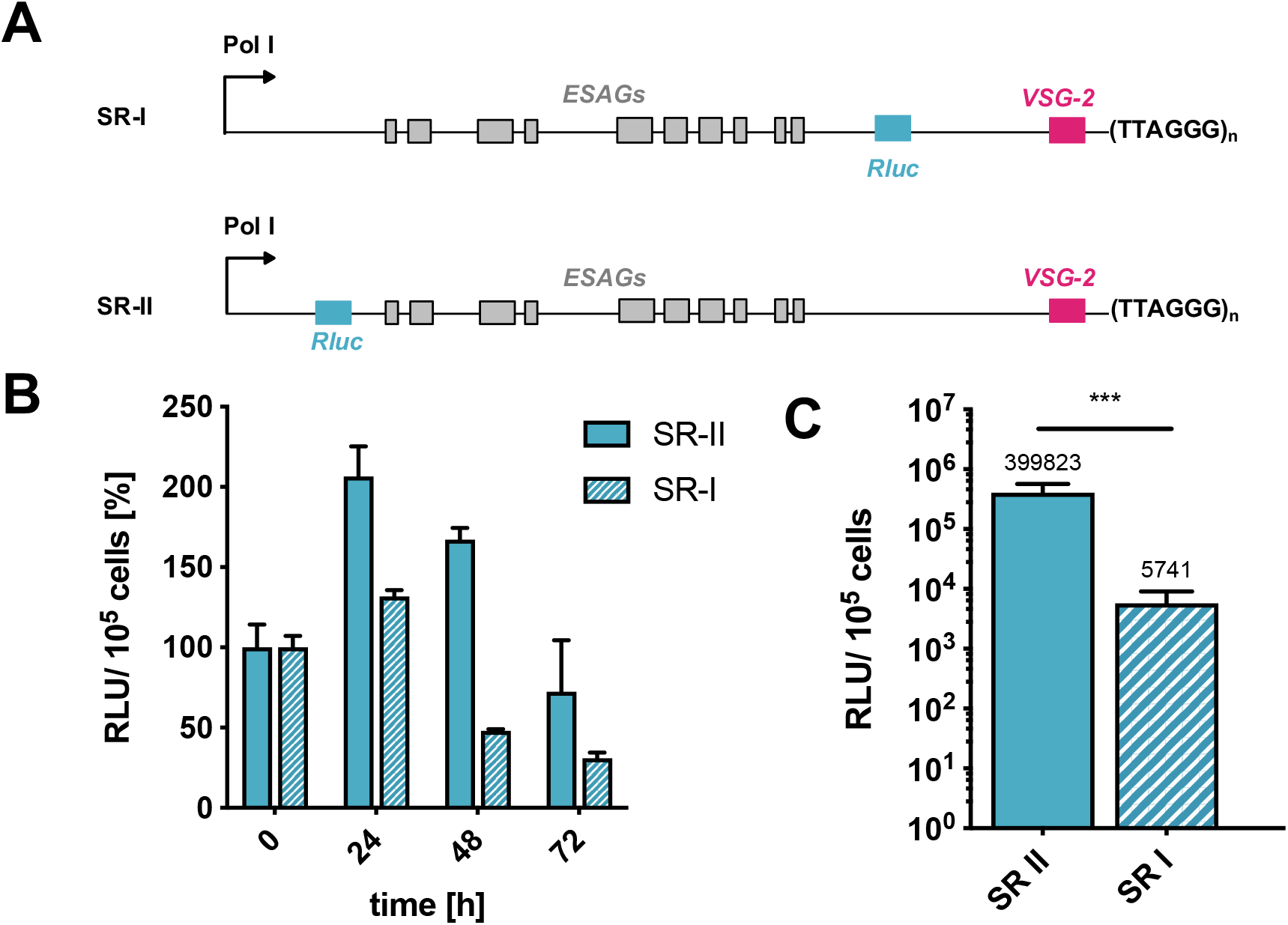
Faster silencing kinetics at the telomere region of the active BES. BES silencing during differentiation of BSF cells to PCF cells. (**A**) Illustration of single Renilla luciferase reporter cell lines with the luciferase either at the telomere (SR-I) or promoter (SR-II) region. (**B**) The Rluc activity was measured at different timepoints upon induction of differentiation. Relative light units (RLU) are presented in percentage, timepoint 0 h was set as 100%. Error bars represent standard deviation of n=3. (**C**) Luciferase activity measurement in BSF cells revealed higher Renilla activity at the promoter region compared to the telomere region. Luciferase activity is shown in relative light units (RLU) at a logarithmic scale. Error bars represent standard deviation of n=15. Mean value is indicated above each column. Statistical significance is shown by an unpaired t-test (***P < 0.001).

**Fig 2:**
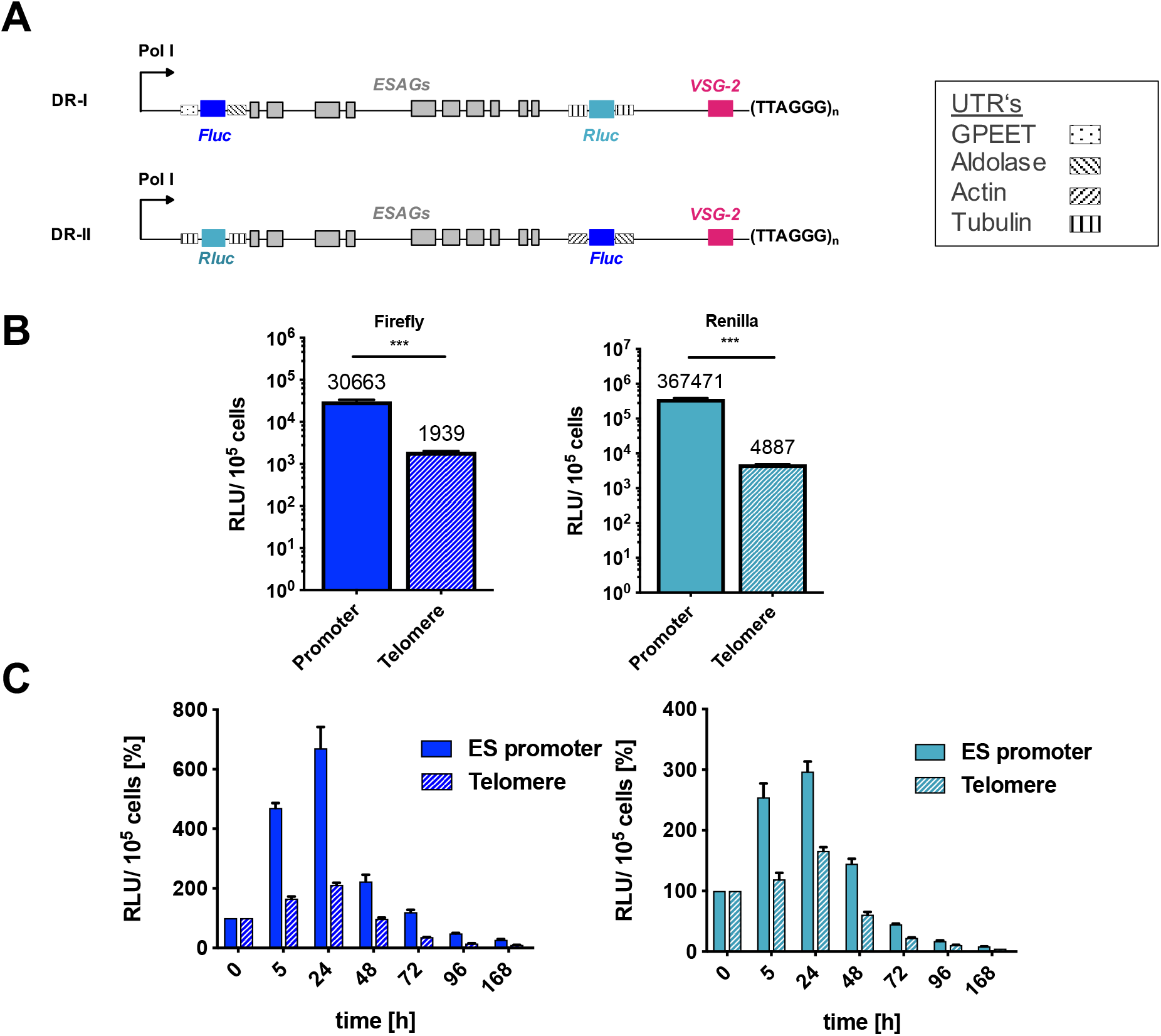
Higher transcriptional activity at the promoter region of the active BES. (**A**) Illustration of the dual luciferase reporter system. The active BES was marked with a Firefly reporter downstream of the BES promoter and a Renilla reporter upstream of VSG 221 (DR-I) and vice versa (DR-II). The UTRs of the Luciferase reporters are marked and described in the associated box on the right (**B**) Luciferase activity of DR-I and DR-II was higher at the promoter region of the active VSG 221 BES than at the telomere. Luciferase activity is shown in relative light units (RLU) at a logarithmic scale. Mean value is indicated above each column. (**C**) Analysis of luciferase activity upon induction of differentiation showed faster silencing kinetics at the telomere region, comparable and already observed in the SR cell lines. Luciferase activity is shown in relative light units (RLU) as absolute values. Error bars represent standard deviation of n=3. Statistical significance is shown by an unpaired t-test (***P < 0.001).

**Fig 3:**
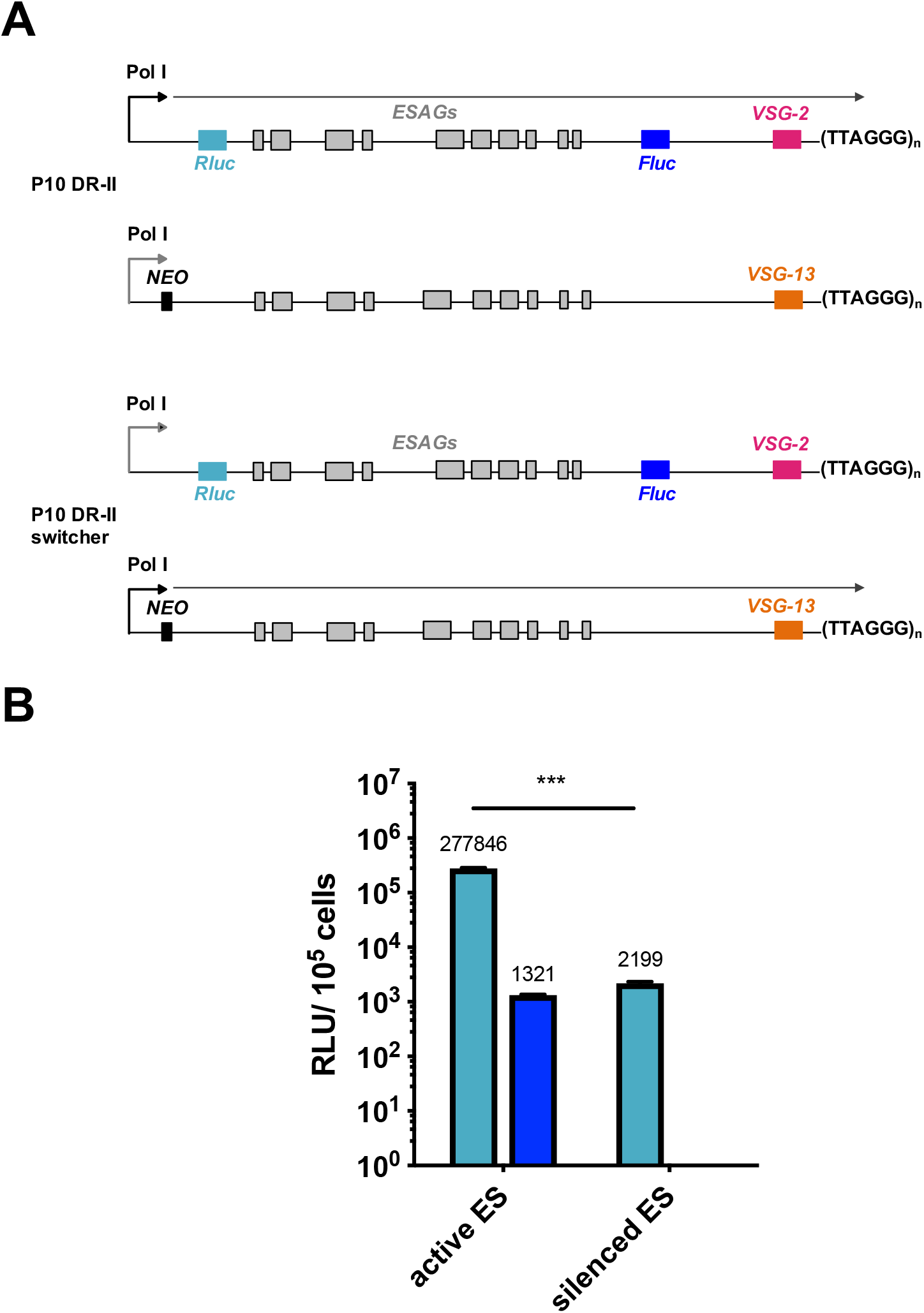
Luciferase activity at active vs. inactive BES. (**A**) Luciferase assays were conducted with a P10-DR II cell line containing Rluc downstream of the BES promoter and Fluc upstream of the active VSG-2 BES (BES1). BES 17 contains an aminoglycoside phosphotransferase resistance gene (NEO). By antibiotic selection with G418, selection for in situ switchers to an active VSG-13 BES (BES17) could be obtained, containing the luciferase reporters in the inactive BES1. (**B**) Measurement of the luciferase reporter activity in P10 DR-II and P10 DR-II switcher. Luciferase activity is shown in relative light units (RLU) at a logarithmic scale. Error bars represent standard deviation of n=3. Mean value is indicated above each column. Statistical significance is shown by an unpaired t-test (***P < 0.001, n=3).

**Fig 4:**
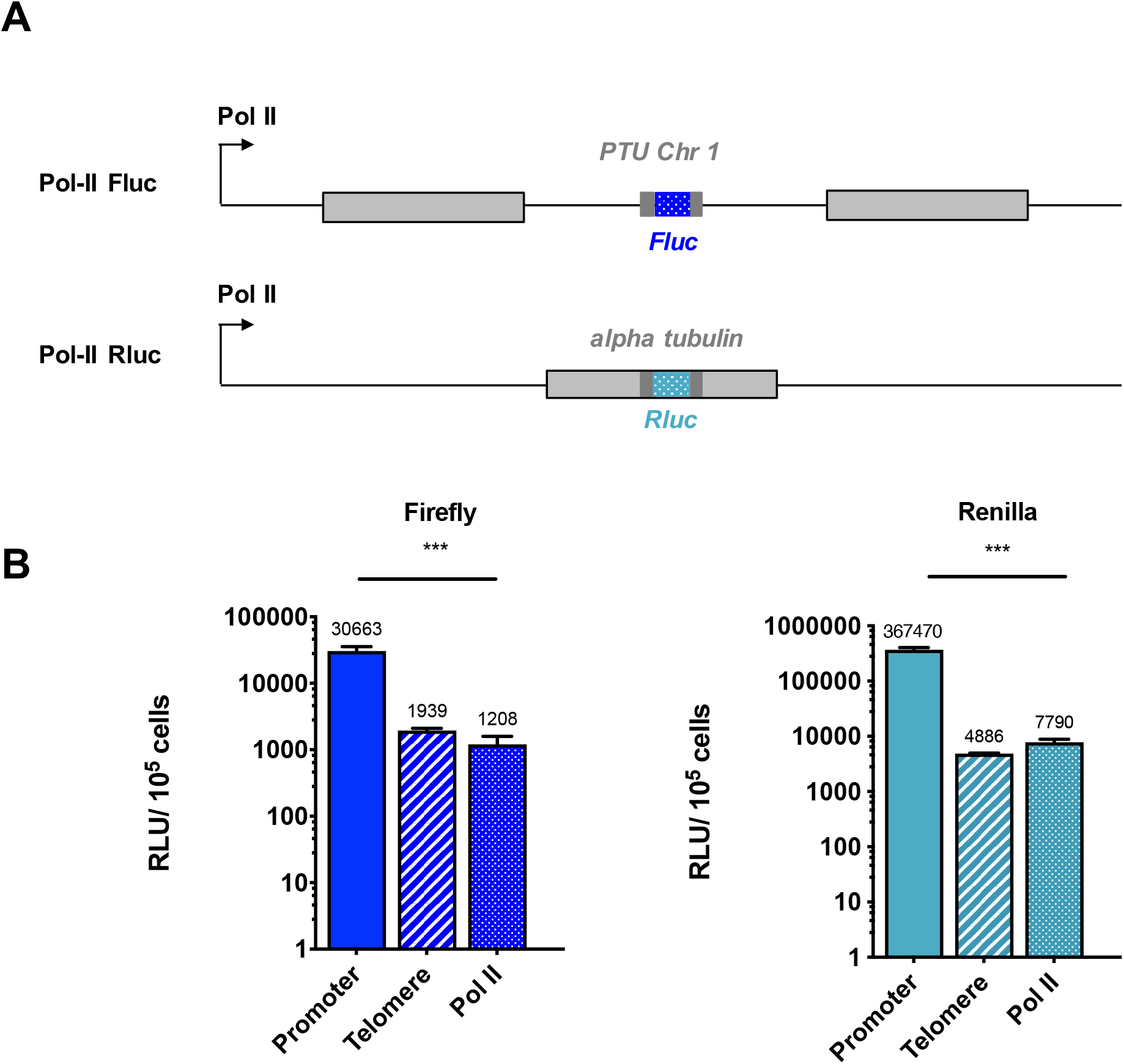
Luciferase activity at the telomere of the active BES is comparable to a Pol-II-transcribed locus. (**A**) Single luciferase reporters were integrated within a PTU of Chromosome 1 (Pol-II Fluc) or at the alpha tubulin region (Pol-II Rluc). (**B**) The Fluc activity of DR-II (Fig 2A) was comparable to the activity of Pol-II Fluc; the Rluc activity of DR-I (Fig. 2A) was comparable to Pol-II Rluc. Luciferase activity is shown in relative light units (RLU) on a logarithmic scale. Error bars represent standard deviation of n=3. Mean value is indicated above each column. Statistical significance is shown by an unpaired t-test (***P < 0.001).

## DISCUSSION

Previous studies already described that the transcriptional activity of the active BES is considerably higher compared to the silent BES, but that the silent BES is still expressed in a transcriptionally attenuated manner [48, 49]. 20% of the ESAG6 mRNA is still transcribed from the silent BES [50]. Horn and Cross showed that the expression of a bleomycin resistance protein gene integrated proximal to VSG-2 is over 1000-fold lower when BES1 is silent instead of active [51]. If a reporter gene was integrated directly downstream of the BES promoter, developmental silencing of the active BES led to a transcriptional reduction between approximately 100- [52, 53] and 1000-fold [31, 46]. Transcriptional attenuation of silent BES appears to be the major mechanism to guarantee monoallelic expression of VSG genes. Our study suggests that a similar mechanism is also operating at the active BES. We observed up to 70-fold higher activity of luciferase reporters when they were positioned close to the promoter of the active BES instead of near the telomere. The reporter signals from a telomere-proximal locus in an active BES were comparable to the signals of reporters integrated into loci transcribed by RNA Pol II, indicating that the transcriptional activity of Pol I near the telomere is similar to the transcriptional activity of Pol II in PTUs. This is consistent with a high rate of transcriptional initiation, but impaired elongation along the active BES. We propose that high Pol I-mediated transcriptional activity is needed to counteract inefficient transcriptional elongation to guarantee sufficient transcription of the VSG gene itself at the end of the BES. It is not intuitive that transcription is attenuated at an active BES considering that substantial amounts of VSG is needed to cover the parasite’s surface. However, this might be a tradeoff to balance between different molecular mechanisms that have to guarantee sufficient transcriptional activity at the VSG gene, thigh monoallelic transcription, the capacity for VSG gene switching and developmental regulation of the BES.

Why hasn’t this been observed before? Most reports about BES transcriptional regulation focus on the difference between active and silent BES [46], BES activity during developmental differentiation, or in PCFs [54]. In most studies, only one reporter gene targeting one locus was integrated. When integrating reporters at different positions within the BES, a promoter was often placed in front of the reporter gene, circumventing endogenous expression of the BES, or the reporter genes were located within several kilobase pairs distance to the telomeres [55]. This is the first time Pol I activity under the control of an endogenous BES promoter at the active BES was systematically monitored with comparable reporter genes integrating at different loci within the same BES.

It has long been assumed that the high Pol I processivity is needed for strong VSG gene transcription. Our data suggests a different scenario. We suggest that high Pol I activity is needed to allow a moderate activity at the end of an attenuated BES. There are other important molecular mechanisms, which guarantee for high VSG protein levels needed to cover the parasites surface. Post-transcriptional factors influencing mRNA abundance are already known to play a significant role [56, 57]. The untranslated regions of the mRNA have an effect on trans-splicing and 3’-polyadenylation, affecting mRNA stability [58]. RNA-binding proteins can recognize specific elements of the 3’UTR and influence RNA processing, transport and stability as well as translation of the mRNA [59]. Several of these proteins are also involved in regulation of gene expression during different developmental stages during the life cycle of *T. brucei [60]*. During the differentiation process into the procyclic form, the VSG mRNA becomes unstable [61]. In contrast, the half-life of VSG mRNA in BSF is 4.5 h, which is very high compared to the average of a few minutes for other mRNAs [61, 62]. VSG mRNA is therefore the most abundant mRNA in *T. brucei*. It is not fully understood how VSG mRNA stability is regulated. One factor already known to influence the high abundance of the VSG mRNA is the 16-mer motif of the 3’UTR [63]. Viegas et al. propose that inclusion of N^6^-methyladenosine in the poly(A) tails of VSG stabilizes VSG transcripts [64]. Very recently, Faria *et al* reported that the active VSG gene is always associated with the splice leader RNA array leading to efficient VSG mRNA processing [65]. This might also be an important mechanism to compensate for attenuated transcriptional activity at the VSG gene. Additionally, the VSG protein itself is also very stable. Although turnover of the VSG cell-surface pool happens within approximately 12 minutes [66], it is manly mediated by rapid endocytosis and recycling of the protein and VSG shedding into the environment [67]. The half-life of the VSG protein itself has been estimated to be approximately t^1^/_2_ = 237 h ^+^/_-_ 45 h hours in BSF [68]. These mechanisms can all contribute to fulfill the need for the high demand of VSG proteins on the parasite surface without the need of high transcriptional activity at VSG genes, which might be harmful for the parasite. A highly active polymerase at the VSG gene might cause read-through transcription from the telomeric repeats that start directly down-stream of the VSG gene. However, telomeric transcripts form RNA:DNA hybrids, which are toxic for the cell [69]. Hence, the interplay between transcriptional attenuation and a molecular machinery that compensates for low transcription rates at the VSG gene might be a trade-off between the need for a highly abundant VSG mRNA and genomic stability.

## ACKNOWLEDGEMENTS

We thank Brooke Morriswood and Lucy Glover for carefully reading this manuscript and for all comments and suggestions. Work in the Janzen laboratory was supported by a research grant of the Deutsche Forschungsgemeinschaft (JA1013/6-1).

## Supplementary Figures

**Fig S1:**
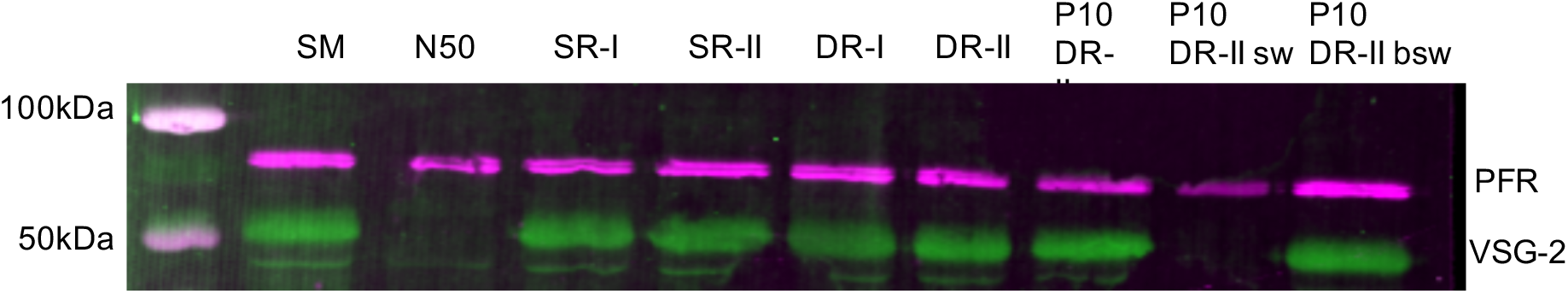
VSG-2 expression in reporter cell lines. Western blot analysis of VSG-2 expression in parental cells (SM) and reporter cell lines. N50 serves as negative control for VSG-2 expression. VSG-2 expression was monitored with an anti VSG-2 antibody. PFR1,2 proteins were detected with anti-PFR1,2 antibody and were used as a loading control.

**Fig S2:**
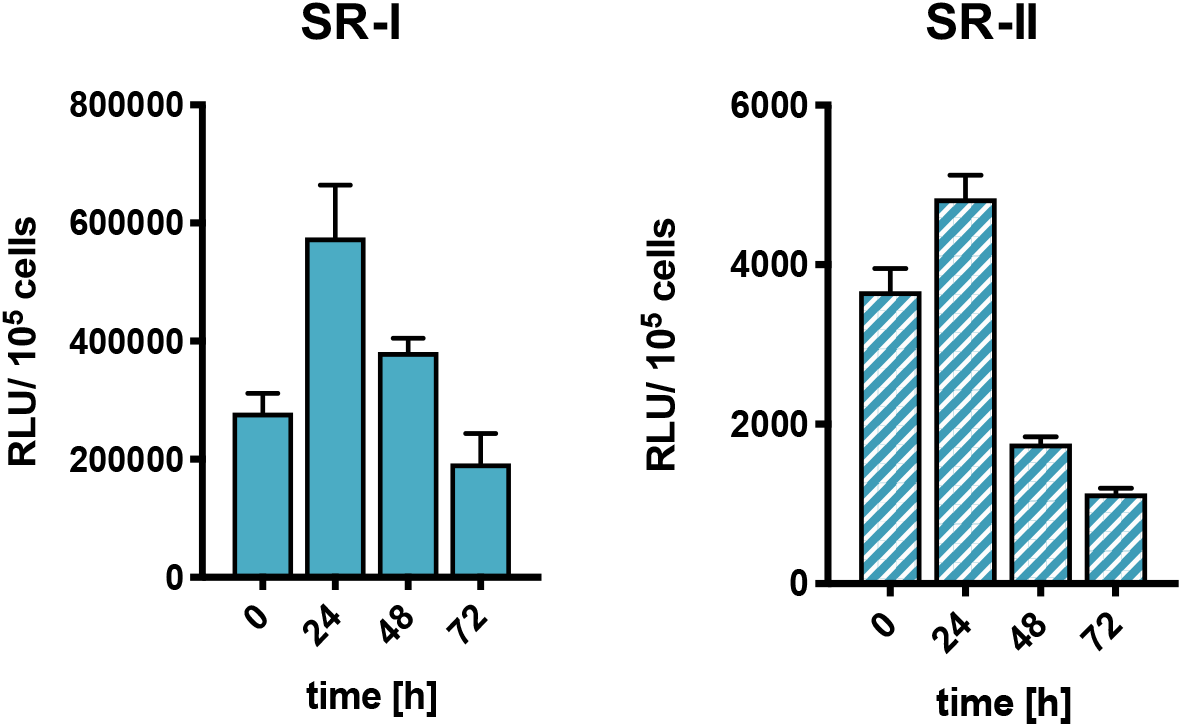
Analysis of the Rluc activity of single reporter cell lines during differentiation from BSF to PCF. Cell lines contained a Renilla luciferase reporter at the BES promoter (SR-II) or at the telomere region (SR-I). Measurement occurred at different time points after induction of differentiation. Luciferase activity is shown in relative light units (RLU) presented in absolute values. Error bars represent a standard deviation of n=3.

**Fig S3:**
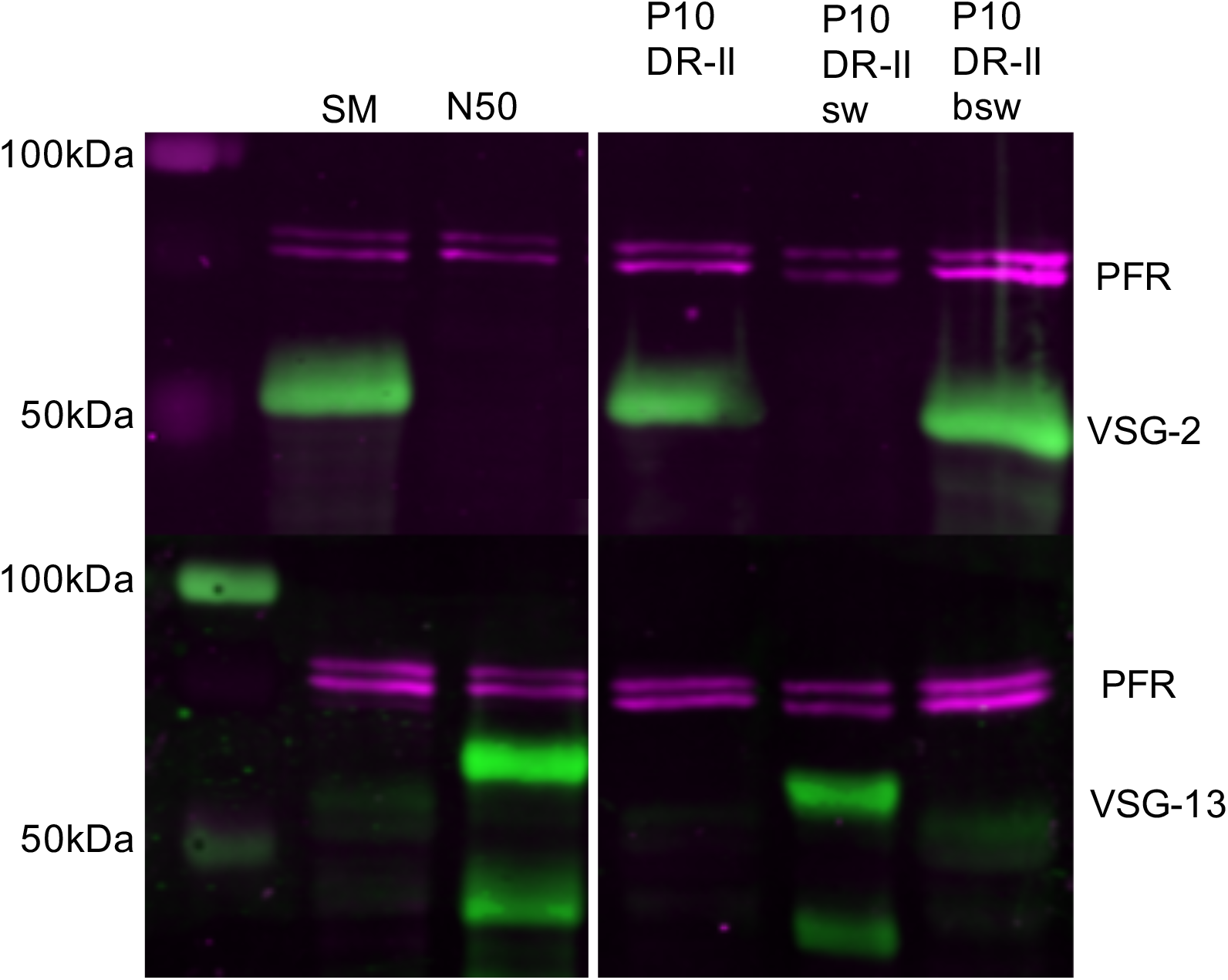
Western blot analysis of P10 DR-II cell lines with CRD-depleted anti-VSG-2 and anti-VSG-13 antibodies to confirm successful switching. P10 DR-II and P10 DR-II “ back switcher” (bsw) expressed VSG-2, P10 DR-II switcher (sw) expressed VSG-13. SM and N50 whole cell lysates served as control for either VSG-2 or VSG-13 expression. PFR protein expression detected with anti-PFR antibody was used as loading control.

